# Fluoxetine for Prenatal Alcohol-Exposed Mice: Addressing Impaired Enrichment-Mediated Neurogenesis

**DOI:** 10.64898/2026.03.13.711456

**Authors:** Arasely M. Rodriguez, Korbin C. Bauer, Elif Tunc-Ozcan, Lee Anna Cunningham

## Abstract

**Background:** Fetal alcohol spectrum disorder (FASD) encompasses a variety of disorders that occur after a fetus has been exposed to alcohol. Hippocampal related issues are a common neurological deficit found in FASD. The dentate gyrus within the hippocampus is a unique area of the brain that continues to generate new neurons into adulthood. This neurogenesis can be enhanced by an enriched environment (EE); however in prenatal alcohol exposed (PAE) mice this EE-mediated neurogenesis is impaired. In addition to EE, selective serotonin reuptake inhibitors (SSRIs), such as fluoxetine, also promote neurogenesis. Here we examine if fluoxetine restores the impaired EE-mediated neurogenesis of PAE mice.

**Method:** PAE mice were generated using a voluntary limited access model where mice received a 10% ethanol (w/v) solution during gestation. To evaluate neurogenesis, we use a NestinCre^T2^:tdTomato transgenic mouse line in which newborn dentate granule cells (nDGCs) can be evaluated by tdTomato flourescence. PAE and saccharine control (SAC) mice were placed in either standard housing (SH) or enriched environment (EE). Subsequently, we administered fluoxetine (FLX) or vehicle (VEH) after which neurogenesis was evaluated.

**Results:** PAE resulted in impaired EE-mediated neurogenesis. This neurogenic impairment was not restored by FLX. Interestingly, FLX did increase neurogenesis in PAE mice while housed in SH.

**Conclusion:** These results suggest that there is a neurogenic ceiling in PAE mice that cannot be increased by fluoxetine in EE. However, fluoxetine can increase neurogenesis while the environment is less complex.

## Introduction

Alcohol consumption during pregnancy remains an ongoing public health concern. Studies have estimated fetal alcohol spectrum disorders (FASDs) in 1 to 5 per 100 school-aged children in the United States (May *et al*., 2018, 2014, 2009). Fetal alcohol syndrome (FAS), the more severe form of FASD, cost the US $156.7 million to $8.5 billion (Popova *et al*., 2021). As this estimate only considers the more easily diagnosable FAS, the true economic cost of FASD is likely higher. Rates of women who have reported drinking during pregnancy have increased (CDC, 2022). 13.5% of women who drank during pregnancy reported that they were still drinking, and 5% responded that they had engaged in binge drinking within the last 30 days (CDC, 2022). With increases in drinking rates in women who are pregnant, there is a need for potential therapeutic options for people with FASD, as even moderate levels of alcohol exposure during gestation can result in cognitive impairments in children (Valenzuela *et al*., 2012).

FASD has long-lasting effects on many brain regions, with hippocampal deficits well-documented in clinical FASD and in preclinical models (review Cunningham et al., 2025; Caputo et al., 2016). For example, MRI studies of children with FASD have shown reduced hippocampal volumes, which are correlated with learning and memory deficits (Norman *et al*., 2009). Additionally, FASD patients are commonly diagnosed with depression and anxiety, mental illnesses that have been related to hippocampal function (review Mattson et al., 2019). Furthermore, preclinical rodent models of FASD have documented long-lasting impairment in enriched environment (EE)-mediated adult hippocampal neurogenesis, which involves the production of new neurons in the dentate gyrus of the adult hippocampus (review Gil-Mohapel et al., 2010).

Adult hippocampal neurogenesis is the generation of newborn dentate granule cells (nDGCs) from a pool of progenitor cells that persist within the dentate gyrus of the hippocampus throughout life. Previous work has demonstrated that PAE mice exposed to moderate levels of alcohol throughout gestation display an ~50% impairment of EE-mediated adult hippocampal neurogenesis compared to control mice in EE (Gustus et al., 2020; Kajimoto et al., 2016, 2013; Choi et al., 2005). This impairment is also associated with impaired learning, impaired dendritic plasticity, and altered activation of dentate granule neurons. In the present study, we tested whether treatment with fluoxetine can reverse the neurogenic deficit in PAE mice exposed to EE.

Fluoxetine, an antidepressant and a selective serotonin reuptake inhibitor (SSRI), has been routinely used to increase neurogenesis in preclinical models of depression. In fact, the increase of adult hippocampal neurogenesis by antidepressants, including fluoxetine, has been suggested as one of the therapeutic mechanisms of antidepressants (Planchez, Surget and Belzung, 2020). As depression and anxiety are the most common mental health diagnoses in patients with FASD, it is not surprising that SSRIs are the 3^rd^ and 5^th^ most commonly prescribed in this population, depending on Medicaid or private insurance coverage, respectively (Senturias, Ali and West, 2022). However, studies on the efficacy of antidepressants in these patients remain lacking. In fact, a review by Ritfeld et al. (2022) found that the only study to measure the efficacy of antidepressant in patients with FASD was conducted by Coe et al. (2001). Coe et al. found that children with FASD responded positively to sertraline, an SSRI.

Here we are investigating fluoxetine, a neurogenic antidepressant, as a potential therapeutic intervention for impaired EE-mediated hippocampal neurogenesis in PAE mice. We used a NestinCre^T2^:tdTomato PAE mice subjected to a limited access, drinking-in-the-dark PAE paradigm. These PAE mice were housed in standard housing (SH) or EE conditions. In their respective housing, mice received fluoxetine (FLX) or water as vehicle control (VEH). We measured depression-like and anxiety-like behavior; quantified neurogenesis through fluorescently tagged newborn neurons; and measured serotonin fibers.

## Methods and Materials

### Mice

Nestin-CreER^T2^:tdTomato transgenic mice were used for these studies. These mice were obtained from breeding colonies maintained in the University of New Mexico Health Sciences Center Animal Resource Facility. Colonies were maintained for homozygosity at both the Nestin-CreER^T2^ (Lagace *et al*., 2007) and Ai9 (RCL-tdT) bitransgenic gene loci (Madisen *et al*., 2010), on a C57BL/6J background. Genotyping was routinely performed on tail tissue by qualitative PCR services performed by TransnetYX genotyping (Cordova, TN), using previously published primer sequences (Gustus *et al*., 2019). Mice were housed under reverse 12-hr dark/12-hr light cycle (light off at 08:00h) in a humidity and temperature-controlled room with *ad libitum* food (Teklad 2920X) and water. Animal procedures were approved by the University of New Mexico Institutional Animal Care and Use Committee in accordance with NIH policies on Human Care and Use of Laboratory Animals.

### Limited Access Drinking in the Dark Prenatal Alcohol Exposure

PAE offspring were generated within the New Mexico Alcohol Research Center Scientific Core using a 1-2^nd^ trimester human equivalent mouse, limited-access, drinking-in-the-dark alcohol exposure paradigm as characterized by Brady et al. (2012). Dams are transferred to a room with minimal sound interruption and acclimate to single housing for a week. After acclimatization, two hours into their awake phase dams are initially provided 5% ethanol (Decon Labs, King of Prussia, PA, USA) in 0.066% SAC (Sigma-Aldrich, St. Louis, MO, USA) or 0.04% saccharin in water as control (SAC) for 4 hrs. After four days the ethanol concentration was then increased to 10%, so that ethanol drinking mice received 10% ethanol in 0.066% SAC instead of drinking water for 4 hrs. Mice that only received 0.04% SAC in water served as a control. After drinking at 10% for 2 weeks, mice were bred by placing the females into the male’s cage for 2 hrs, immediately following the 4 hr drinking period. Breeding was repeated for 5 consecutive days. Dams continued to drink alcohol throughout the gestation period. After parturition, ethanol drinking was ramped down in concentrations of 5%--2.5%--0% with bouts of 2 days each. Blood alcohol levels (BACs; mg/dL) during the 4 hr access period, as determined for dams after two weeks of drinking 10% alcohol, was 40.55 +/− 29.46, n = 10. Offspring were weaned and separated by sex at postnatal day 24 and left undisturbed except regular care until tamoxifen injections.

### Drug Administration

Tamoxifen (Sigma-Aldrich) was dissolved in filter sterilized sunflower oil (Sigma-Aldrich; 30 mg/mL) via a heated water bath sonicator. All mice (postnatal 40-50) were administered tamoxifen for a period of 5 consecutive days via intra peritoneal injection (i.p) at a dose of 120 mg/kg, to visualize and quantify nDCGs as previously described by Gustus et al. (2020). FLX (Tokyo Chemical Industry, Portland, OR) was dissolved in bacteriostatic water (Pfizer, New York, NY) at a concentration of 2 mg/mL. FLX was administered via i.p. for 3 weeks at a dose of 10 mg/kg, as previously described by Tunc-Ozcan et al. (2021), and mice injected with water served as FLX controls.

### Animal Housing

After tamoxifen administration, mice were placed in either SH or EE. EE housing consisted of a large cage (48 cm x 27 cm x 20 cm, Type IV) with 6-8 mice per cage as previously described (Choi, Allan and Cunningham, 2005; Kajimoto, Allan and Cunningham, 2013; Kajimoto *et al*., 2016; Gustus *et al*., 2020). The cage had a total of seven enrichment items: two running wheels, two tunnels, a wooden ladder, a rodent house, and a dangling toy. Note, every week the toys were swapped with clean ones and placed in a different area in the cage. For SH, mice were placed in a standard cage (28 cm x 18 cm x 13 cm) with a single disposable rodent house, 2-4 mice per cage.

### Behavior Testing

Before behavioral testing, mice were acclimated to the testing room for a period of 2 hrs. The open field test (OFT) was conducted in a white square arena (30 × 30 cm). Light was adjusted so that the corners of the arena were ~45 lux and the center of the box was at ~90 lux. The box was disinfected with 70% ethanol and mice were placed in the center of the box and recorded for a total of 8 min. The day after OFT testing, mice were run through the tail suspension test (TST). The TST arena was set up under ~90 lux. A 4 cm cylinder was placed on the mouse’s tail to prevent tail climbing. A piece of 7 cm tape was placed on the tail with 1 cm overlap around the tail. Mice were then suspended from their taped tails for a duration of 5 min. Behavioral quantification for OFT and TST was conducted with Ethovision XT 17 (Noldus, Leesburg, VA) software.

### Quantification of tdTomato positive cells

After sodium pentobarbital (Zoetis, Florham Park, NJ) overdose (i.p;150 mg/kg) mice were transcardially perfused with 0.1% procaine (Sigma-Aldrich) and 2 U/mL heparin (Fresenius Kabi USA, Lake Zurich, IL) in phosphate buffered saline (PBS), then with a 4% paraformaldehyde (PFA; Sigma-Aldrich) solution in phosphate buffer. Brains were kept in 4% PFA for 24hrs and subsequently dehydrated in 30% sucrose in PBS. Coronal hippocampal sections of 40 um were taken from AP −1.755:−2.48 mm bregma. Sections were coverslipped with 1.5 oz coverglass with DAPI Flourmount G media (Electron Microscopy Sciences, Hatfield, PA). Images were taken at 20x on Zeiss Axioscan 7. tdTomato positive cells within the superior blade of the dentate gyrus were manually counted using QuPath v0.5.1 (Bankhead *et al*., 2017) software. The average of three sections per mouse was taken as tdTomato+ value.

### Quantification of SERT Fibers

Fixed sections, 40 μm thick, were washed in PBS for 3 × 5 min each. Sections were then permeabilized in 0.04% Triton X for 10 min, followed by 3 × 5 min washes in PBS. Following this, sections were incubated in Image IT FX for 30 min and washed 1x in PBS for 5 min. After that, sections were blocked in 0.25% Triton X 100 with 10% Donkey Serum in PBS for 1 hr and incubated in primary antibody (HTT/SERT pAB GP; MSFR103300; AB_2571777; Nittobo Medical; 1:500) in 0.25% Triton X 100 + 1 % BSA in PBS for 24 hr at room temperature. Sections were then washed in PBS for 3 × 5 min and incubated with secondary antibody (Alexa Flour 488 AffiniPure Goat Anti Guinea Pig IgG; 106-545-003; AB_2337438; Jackson ImmunoResearch; 1:500) for 1 hr, followed by washing 3 × 5 min in PBS. Sections were mounted and imaged on a Zeiss LSM800 confocal microscope at 20X. SERT fibers were quantified using QuPath v0.5.1 (Bankhead et al. 2017). Pixel Classification with a random trees model trained on 3 representative images with 0.62 um/px resolution. ROIs were defined in the molecular layer, above the superior blade of the dentate gyrus.

### Statistical Analysis

This study included male and female offspring. A complete litter was removed from our data analysis due to failed CRE-mediated recombination, likely due to experimental error in genotyping. Number of mice found in Sup Table 1. All remaining data were analyzed as follows and indicated in figures: using one-way t-test to facilitate *a-priori* comparisons in neurogenesis, two-way t-test to facilitate *a-priori* comparisons in SERT fibers and behavior; and welch’s t-test (denoted with t^w^) for data with unequal variances when there were unequal variances. Analysis was conducted on GraphPad Prism v10.1 (La Jolla, CA). Data are expressed as mean ± SEM to two significant figures, with p-values <0.05 considered significant. Error bars are SEM.

## Results

### FLX increased adult neurogenesis in both SAC and PAE mice under SH conditions

We first evaluated FLX efficacy to stimulate neurogenesis in mice housed under SH. We counted the number of tdTomato positive cells across 3 hippocampal sections in each mouse, as a measure of neurogenesis. As shown in Fig. 2B, FLX significantly increased neurogenesis (t(13) = 1.9, p = 0.0368) in SH-SAC-FLX (63 +/− 5.2) compared to SH-SAC-VEH (46 +/− 6.5). FLX also significantly increased neurogenesis (t(21) = 3.5, p = 0.0010) in SH-PAE-FLX (68 +/− 5.8) compared to SH-PAE-VEH (46 +/− 3.5). There was no significant difference in neurogenesis (t(17) = 0.74, p = 0.2340) between SH-SAC-FLX (63 +/− 5.2) and SH-PAE-FLX (68 +/− 5.8). Similarly, baseline neurogenesis was not significantly different (t(17) = 0.10, p = 0.4596) between SH-SAC-VEH (46 +/− 6.5) compared to SH-PAE-VEH (46 +/− 3.5). In conclusion, FLX in SH increased neurogenesis in both SAC and PAE animals.

**Figure 1.**
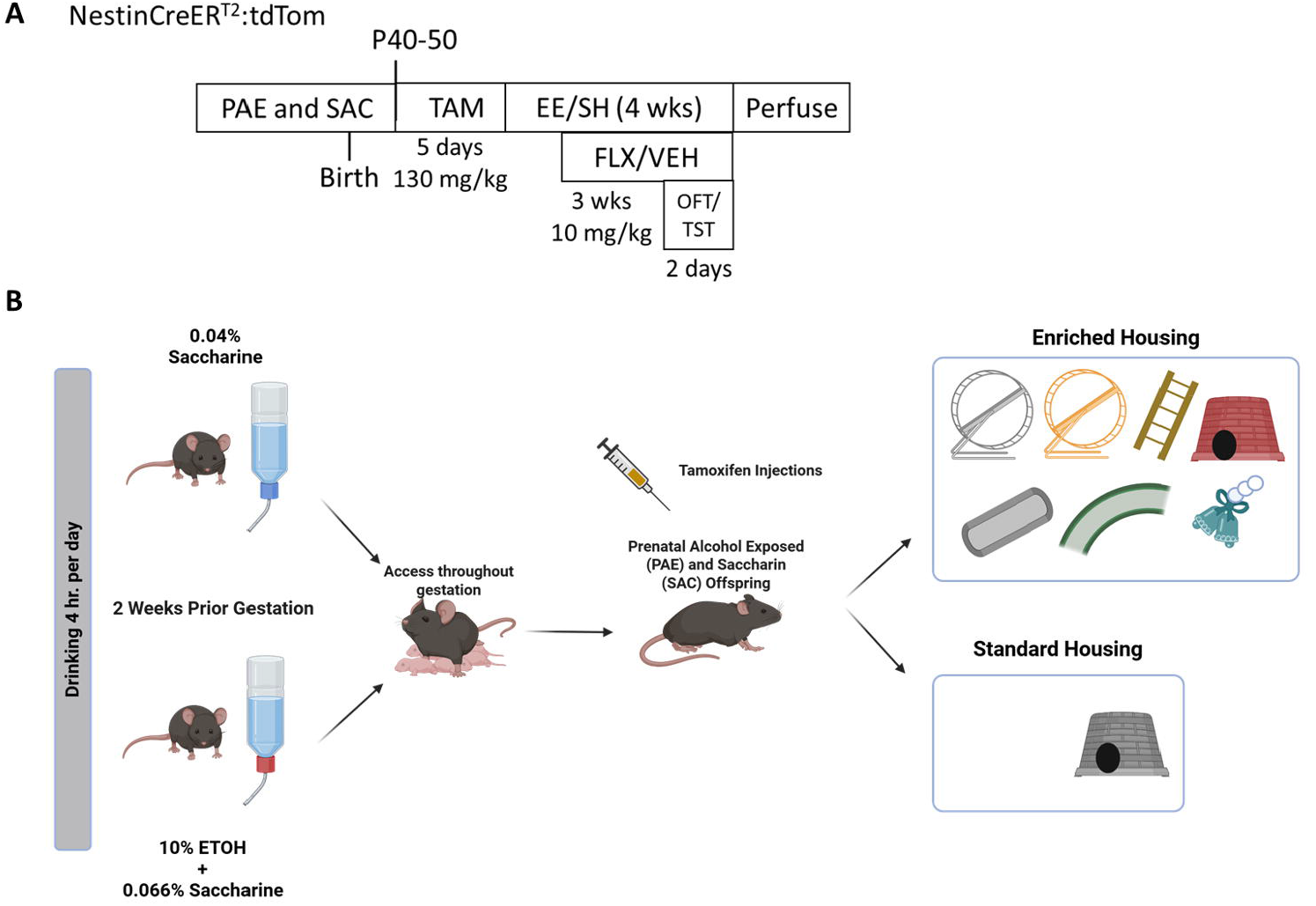
PAE mice and SAC mice generation. A) Experimental timeline. B) Experimental Design.

**Figure. 2.**
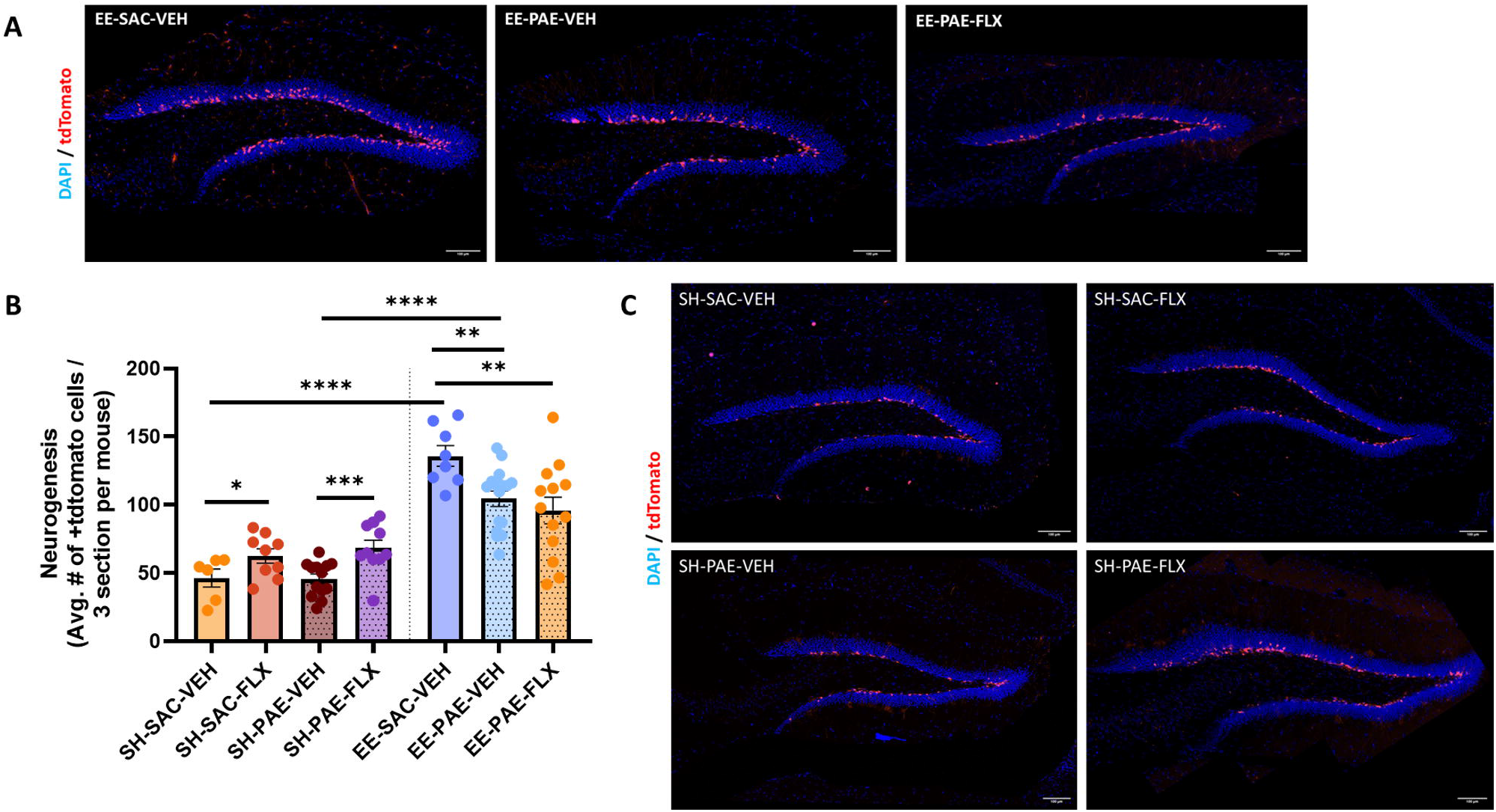
EE-mediated adult neurogenesis in PAE mice is not restored by FLX. A) Representative (20x; scale 100 um) confocal microscopy images of coronal sections through the dorsal dentate gyrus of EE groups demonstrating +tdTomato nDGCs (red), DAPI nuclear counterstain (blue). B) Number of tdTomato+ nDGCs across groups (means ± SEM). SH-SAC-VEH (n = 6); SH-SAC-FLX (n = 9); SH-PAE-VEH (n = 13); SH-PAE-FLX (n = 10); EE-SAC-VEH (n = 8); EE-PAE-VEH (n = 16); EE-PAE-FLX (n = 13). C) Representative confocal microscopy images of coronal sections through the dorsal dentate gyrus of SH groups demonstrating +tdTomato nDGCs (red), DAPI nuclear counterstain (blue).

### EE-mediated adult neurogenesis in PAE mice is not restored by FLX

Having demonstrated that FLX stimulated neurogenesis in both SAC and PAE mice, we next tested whether this FLX administration regimen would restore EE-mediated neurogenesis in PAE mice. As shown in Fig. 2B, EE significantly (t(12) = 8.5, p < 0.0001) increased neurogenesis in EE-SAC-VEH mice (136 +/− 7.6) compared to SH-SAC-VEH (46 +/− 6.5). EE also significantly increased neurogenesis (t(27) = 8.3, p < .0001) in EE-PAE-VEH (104 +/− 5.7) compared to SH-PAE-VEH (46 +/− 3.5). However, neurogenesis was significantly (t(22) = 3.2, p = .0018) higher in EE-SAC-VEH mice (136 +/− 7.6) compared to EE-PAE-VEH (104 +/− 5.7), showing the EE-mediated neurogenic deficit of PAE mice. Neurogenesis in EE-PAE-FLX mice (96 +/− 9.7) was still significantly lower (t(19) = 2.9, p = 0.0047) compared to EE-SAC-FLX (136 +/− 7.6). These results show that deficits seen in neurogenesis as a result of PAE were not reversed by FLX treatment in an EE.

### FLX increases SERT fibers in SAC mice but not in PAE mice under SH conditions

In the same cohort of mice as above, we next went on to determine the effects of FLX on the percent area (μm^2^) of serotonergic nerve fibers. As shown in Fig. 3B, FLX significantly increased (t(9) = 2.5, p = 0.0349) SERT fibers in SH-SAC-FLX (11 +/− 1.5) compared to SH-SAC-VEH (5.2 +/− 1.9). FLX did not significantly increase (t(16) = 0.37, p = 0.7132) SERT fibers in SH-PAE-FLX (7.1 +/− 1.2) compared to SH-PAE-VEH (6.5 +/− 1.1). There SERT fibers were significantly higher in (t(14) = 2.2, p = 0.0488) SH-SAC-FLX (11 +/− 1.5) compared to SH-PAE-FLX (7.1 +/− 1.2). Baseline SERT fibers were not significantly different (t^w^(5.0) = 0.59, p = 0.5790) between SH-SAC-VEH (5.2 +/− 1.9) and SH-PAE-VEH mice (6.5 +/− 1.1). In conclusion, FLX increased serotonin fibers only in SAC mice, while PAE mice did not show this increase.

**Figure. 3.**
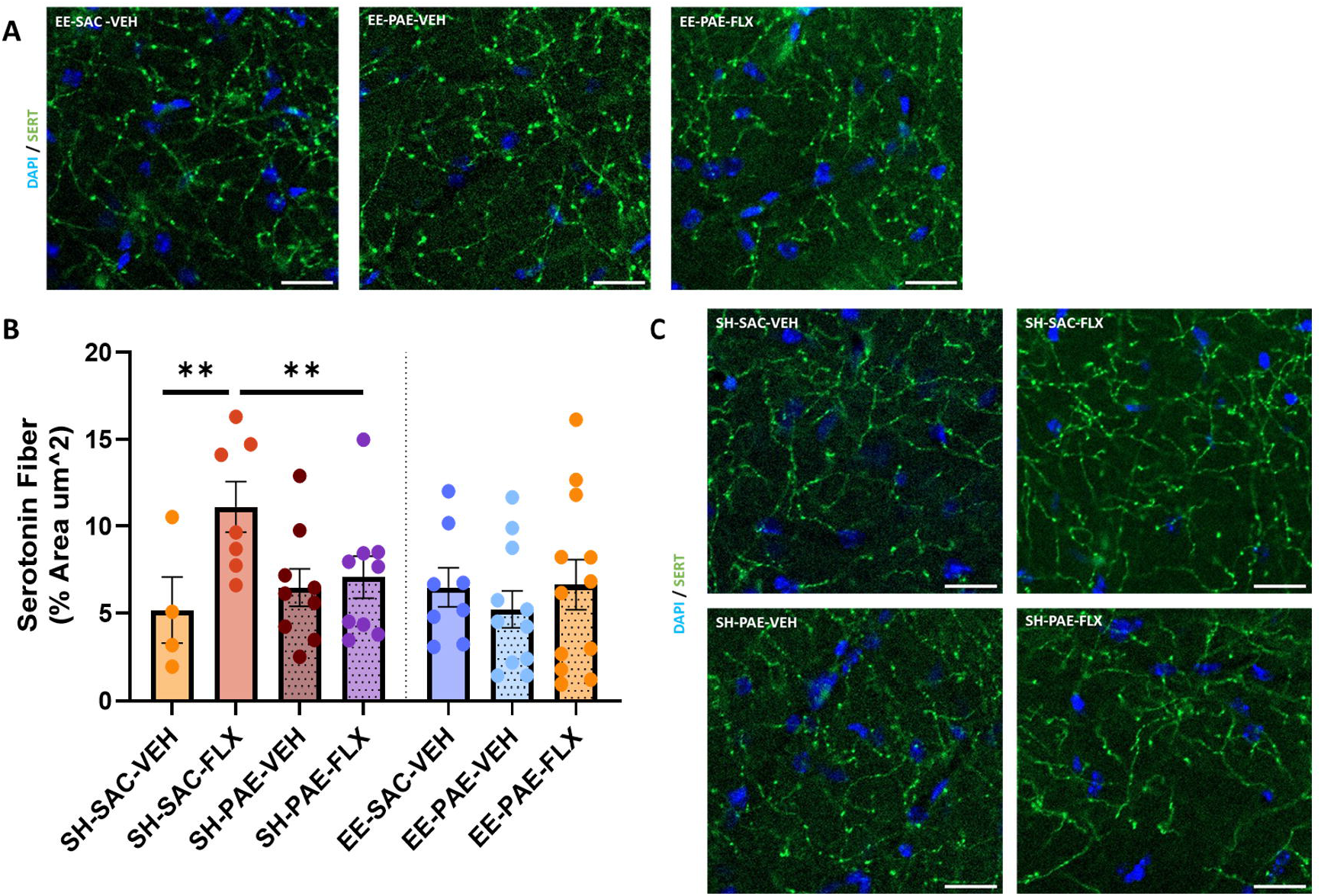
FLX increases SERT fibers in SAC mice but not in PAE mice under SH conditions.. A) Representative (20x; scale 100 um) confocal microscopy images of coronal sections through the molecular zone of the dorsal hippocampus of EE groups demonstrating SERT fibers (green), DAPI nuclear counterstain (blue). B) % Area (um^2^) across groups (means ± SEM). SH-SAC-VEH (n = 4); SH-SAC-FLX (n = 7); SH-PAE-VEH (n = 9); SH-PAE-FLX (n = 9); EE-SAC-VEH (n = 8); EE-PAE-VEH (n = 11); EE-PAE-FLX (n = 12). C) Representative (20x; scale 100 um) confocal microscopy images of coronal sections through the molecular zone of the dorsal hippocampus of SH groups demonstrating SERT fibers (green), DAPI nuclear counterstain (blue).

### EE does not increase SERT fibers in SAC and PAE mice

EE did not significantly alter (t(10) = 0.63, p = 0.5417) SERT fibers in EE-SAC-VEH (6.5 +/− 1.1) compared to SH-SAC-VEH (5.2 +/− 1.9). EE did not significantly alter (t(18) = 0.82, p = 0.4241) SERT fibers in EE-PAE-VEH (5.2 +/− 1.1) compared to SH-PAE-VEH (6.5 +/− 1.1). There was no significant difference in SERT fibers (t(17) = 0.80, p = 0.4333) between EE-SAC-VEH (6.5 +/− 1.1) and EE-PAE-VEH (5.2 +/− 1.1). These results show that housing did not affect serotonin fiber density.

### FLX does not alter anxiety-and depression-like behavior in SAC and PAE mice

We also tested anxiety and depressive-like behaviors prior to sacrifice. To determine if FLX altered anxiety or depression-like behavior, mice were evaluated with the OFT and TST. There was no significant difference in distance spent in center (Fig. 4A; cm) (t(13) = 0.081, p = 0.9364) in SH-SAC-FLX (33 +/− 3.7) compared to SH-SAC-VEH mice (33 +/− 3.3). FLX did not significantly alter the distance spent in center (t(21) = 0.37, p = 0.7154) in SH-PAE-FLX (33 +/− 3.3) compared to SH-PAE-VEH (35 +/− 3.2). There was no significant difference (t(17) = 0.040, p = 0.9684) in the distance spent in center between SH-SAC-FLX (33 +/− 3.7) and SH-PAE-FLX (33 +/− 3.3). Baseline distance spent in center was not significantly different (t(17) = 0.45, p = 0.6615) between SH-SAC-VEH (33 +/− 3.3) and SH-PAE-VEH (35 +/− 3.2). These results show that FLX treatment had no effect on anxiety-like behavior, nor did PAE affect groups as measured by the distance in center in the OFT test.

**Figure 4.**
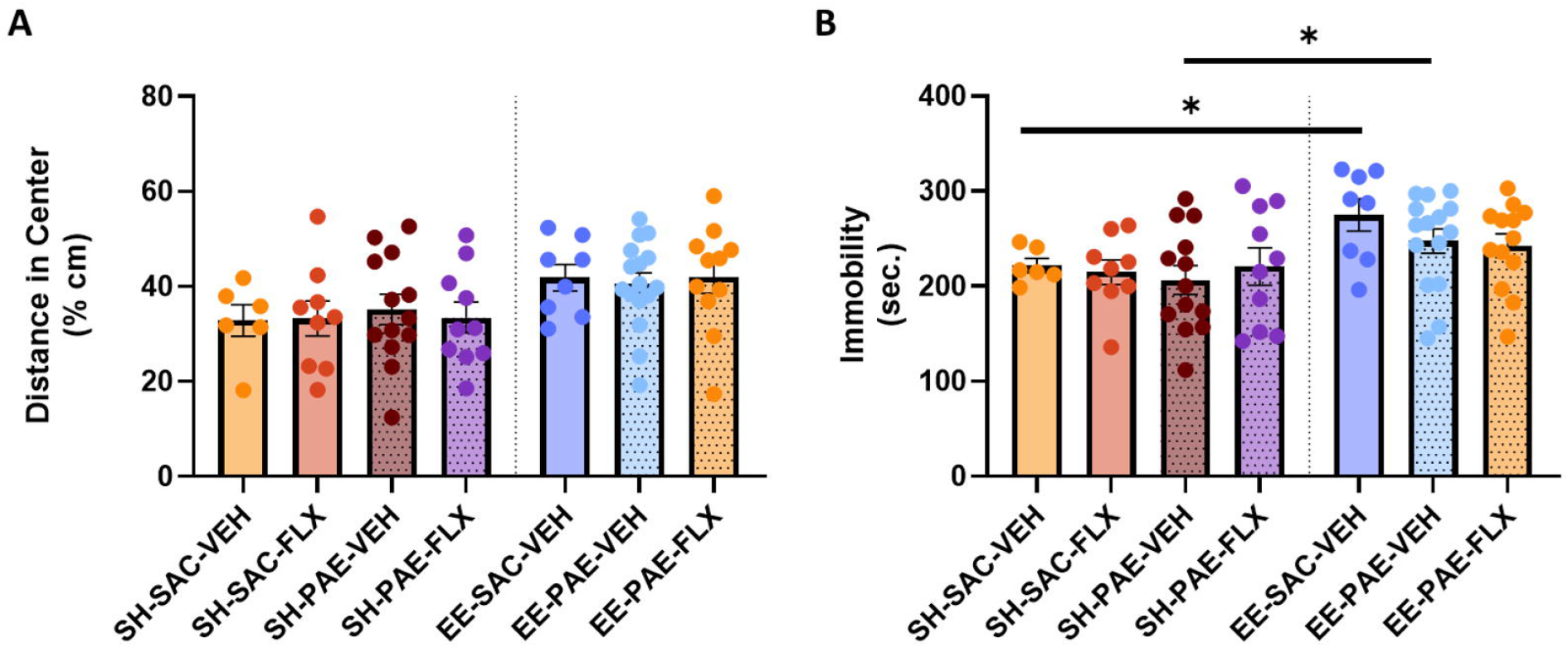
EE increases depression-like behavior in SAC and PAE but does not alter anxiety-like behavior. A) Distance spent in center of OFT arena (% cm) across groups (means +/− SEM). SH-SAC-VEH (n = 6); SH-SAC-FLX (n = 9); SH-PAE-VEH (n = 13); SH-PAE-FLX (n = 10); EE-SAC-VEH (n = 8); EE-PAE-VEH (n = 16); EE-PAE-FLX (n = 11). A) Time spent immobile (sec.) across groups (means +/− SEM). SH-SAC-VEH (n = 6); SH-SAC-FLX (n = 9); SH-PAE-VEH (n = 13); SH-PAE-FLX (n = 10); EE-SAC-VEH (n = 8); EE-PAE-VEH (n = 15); EE-PAE-FLX (n = 13).

There was no significant difference in time spent immobile (Fig. 4B; sec.) (t(13) = 0.40 p = 0.6934) between SH-SAC-FLX (215 +/− 13) and SH-SAC-VEH (222 +/− 7.5). There was no significant difference in time spent immobile (t(21) = 0.58, p = 0.5650) between SH-PAE-FLX (221 +/− 20) and SH-PAE-VEH (206 +/− 15). There was no significant difference in time spent (t(17) = 0.24, p = 0.8116) between SH-SAC-FLX (215 +/− 13) and SH-PAE-FLX (221 +/− 20). Baseline time spent immobile was not significantly different (t^w^(16) = 0.90, p = 0.3793) between SH-SAC-VEH (222 +/− 7.5) and SH-PAE-VEH mice (206 +/− 15). In conclusion, FLX did not affect PAE depression-like behavior.

### EE increases depression-like behavior in SAC and PAE but does not alter anxiety-like behavior

EE did not significantly alter the distance in center (t(12) = 2.1, p = 0.0607) between EE-SAC-VEH (42 +/− 2.8) and SH-SAC-VEH (33 +/− 3.3). There was no significant difference in distance in center (t(27) = 1.4, p = 0.1788) in EE-PAE-VEH (41 +/− 2.3) and SH-PAE-VEH (35 +/− 3.2). There was no significant difference (t(22) = 0.34, p = 0.7356) between EE-SAC-VEH (42 +/− 2.8) and EE-PAE-VEH (41 +/− 2.3). These results show that EE did not have an effect on anxiety-like behavior.

There was a significant increase in time spent immobile (t^w^(9.5) = 2.9, p = 0.0177) in EE-SAC-VEH (275 +/− 17) and SH-SAC-VEH (222 +/− 7.5). There was a significant difference in time spent immobile (t(26) = 2.1, p = 0.0468) in EE-PAE-VEH (247 +/− 13) versus SH-PAE-VEH (206 +/− 15). There was no significant difference (t(21) = 1.3, p = 0.2130) between EE-SAC-VEH (275 +/− 17) and EE-PAE-VEH (247 +/− 13). These results show that EE significantly increased depression like behavior in SAC and PAE mice.

## Discussion

Hippocampal deficits are a well-documented consequence of alcohol consumption during pregnancy. It is known that EE increases neurogenesis and improves hippocampal function (Kempermann, 2019), but previous work in our lab demonstrated that PAE impairs EE-mediated neurogenesis (Gustus et al., 2020; Kajimoto et al., 2013, 2016; Choi et al., 2005). In this present study, we asked whether FLX, an SSRI with known neurogenic effects, could restore EE-mediated neurogenesis in PAE mice. We used Nestin-CreER^T2^:tdTomato mice to evaluate the number of newborn dentate granule cells (nDGCs). Mice were either placed in SH or EE; and treated with VEH or FLX. We replicated our prior finding that EE-mediated neurogenesis is impaired in PAE mice compared to SAC mice. This reduction, however, is not as great as our previous studies (Gustus et al., 2020; Kajimoto et al., 2016, 2013; Choi et al., 2005). This is likely attributable due to lower levels of alcohol exposure than we have seen before (Gustus et al., 2020; Kajimoto et al., 2016, 2013; Choi et al., 2005). Nevertheless, the fact that we saw a reduction in EE-mediated neurogenesis shows that PAE reduces the EE-mediated neurogenic capacity of mice.

To evaluate our main hypothesis, FLX was administered to a group of PAE mice housed in EE conditions. PAE-FLX mice did not show an increase in EE-mediated neurogenesis; in fact, they displayed a similar reduction to PAE mice in EE conditions that received VEH. As observed previously, baseline neurogenesis (SH) remained comparable between SAC and PAE mice (Gustus *et al*., 2020; Kajimoto *et al*., 2016; Kajimoto, Allan and Cunningham, 2013; Choi, Allan and Cunningham, 2005). However, the lack of therapeutic response to FLX in restoring EE-mediated neurogenesis does not reflect resistance to its neurogenic effect in PAE mice. In standard conditions, FLX increased neurogenesis in both SAC and PAE mice, indicating that the response is intact but insufficient to restore impaired EE-mediated neurogenesis in PAE mice.

FLX operates through both serotonin-dependent and serotonin-independent mechanisms. The delay between the modulation of serotonin and therapeutic effects of FLX has led to the identification of additional pathways it influences—most notably, the neurotrophic TrkB/BDNF pathway. This pathway remains active even when serotonergic signaling is suppressed (Levy *et al*., 2019). FLX has been shown to directly bind to TRKB receptors, enhancing BDNF signaling, but importantly, this effect has only been observed following chronic FLX administration (Casarotto *et al*., 2021).This neurotrophic mechanism is particularly relevant given that PAE mice have been found to exhibit alterations in BDNF and other downstream neuroplastic pathways such as NMDA receptor signaling (Brady et al., 2012; Samudio-Ruiz et al., 2010; Caldwell et al., 2008; Sutherland et al., 1998). EE usually increases BDNF and neuroplasticity (review Sale et al., 2014). Our previous work has demonstrated that PAE leads to a reduction in EE-mediated neuroplasticity, as evidenced by decreased dendritic density, reduced cell body size, and fewer dendritic spines (Kajimoto, Allan and Cunningham, 2013; Gustus *et al*., 2020). nDGCs experience a critical period that is driven by NMDAR signaling (Tashiro *et al*., 2006; Åmellem *et al*., 2021). It is during this critical period that we have previously noted that the PAE begin to show reduced number of nDGCs (Kajimoto, Allan and Cunningham, 2013). Taken together, our findings suggest that while FLX-induced neurogenesis remains intact under SH conditions, PAE mice in EEs may encounter a neurogenic ceiling substantially lower than that of SAC mice due to reduced plasticity and thus cell survival. It is important to note that this experiment was conducted without stress-inducing paradigms, indicating that the neurogenic effects of FLX observed here are independent of stress, which is a common variable in many FLX studies (Perez-Caballero *et al*., 2014).

To further explore the serotonergic system integrity, we measured SERT fiber density. We found that FLX increased the serotonin fiber density in SAC mice. This finding contrasts Nazzi et al. (2024) and Nazzi (2019) who found that serotonin fiber quantity decreased in the hippocampus after chronic FLX. This difference can be due to different FLX administration methods, specifically Nazzi et al. used a higher dose of FLX. Potentially, serotonin fiber density may initially increase and subsequently decrease, with this reduction occurring earlier at higher doses. Notably, PAE mice did not show an increase in serotonin fibers when given chronic FLX. At baseline both PAE and SAC mice showed similar levels of serotonin fiber density. This suggests that in PAE mice, a serotonin dependent mechanism does not function normally following FLX treatment. PAE has been shown to reduce the number of serotonin neurons in the dorsal raphe nucleus (Zhou *et al*., 2001). The reduced number of serotonin fibers could be due to a reduced number of serotonin neurons. EE did not have an effect on the serotonin fiber density in PAE and SAC mice. This suggests that EE is not exerting its neurogenic affects through the modulation of the serotonergic system, at least not in a manner detectable by our histological methods.

We next evaluated affective outcomes using open field and tail suspension tests. PAE mice showed similar levels of anxiety- and depression-like behavior. These results, taken together with the fact that we did not add a “stressor” to our experiments, indicate that baseline levels of anxiety- and depression-like behavior are not altered by PAE. However, studies have found that PAE, when combined with acute or chronic stress, shows increases in anxiety- and depression-like behavior above similarly stressed control mice (review Hellemans et al., 2010). FLX did not significantly alter anxiety- or depression-like behaviors in either SAC or PAE mice. It is possible that without the use of stressors, FLX does not alter baseline levels of anxiety- and depression-like behavior. EE did not alter anxiety-like behavior but did increase depression-like behavior in SAC and PAE mice. A study by Karolewicz & Paul (2001) found that social housing, like that employed in our EE, increased immobility in mice. These findings complicate the assumption that EE is uniformly beneficial and raise interesting questions about the paradoxical effects of social enrichment.

This study demonstrates that the neurogenic effects of FLX in PAE mice remain intact in SH mice, but it fails to restore EE-mediated neurogenesis to standard levels, suggesting a plasticity impairment that is not improved by SSRI intervention. In PAE mice, the serotonin system responds differently to FLX, which may be due to differences in the number of serotonin neurons in PAE mice. Affective behaviors in PAE mice under non-stressed conditions did not differ from SAC mice, and FLX did not alter these behaviors in either group. Unexpectedly, EE increased depression-like behavior. This study provides the basis for testing how different environments affect pharmacological effectiveness.

## Acknowledgements

We thank the New Mexico Alcohol Research Center; UNM Comprehensive Cancer Center and the Advanced Light Microscopy Shared Resource; Center for Brain Recovery and Repair and the Pre-Clinical Core. This research was supported by the following grants: NIAAA 5R01AA027462, 1F31AA031920, 2P50AA022534; NCI P30CA118100; NIGMS 5P20GM109089.

